# Measuring robust functional connectivity from resting-state MEG using amplitude and entropy correlation across frequency-bands and temporal scales

**DOI:** 10.1101/2020.03.31.017749

**Authors:** Megan Godfrey, Krish D. Singh

## Abstract

Recent studies have shown how MEG can reveal spatial patterns of functional connectivity using frequency-specific oscillatory coupling measures and that these may be modified in disease. However, there is a need to understand both how repeatable these patterns are across participants and how these measures relate to the moment-to-moment variability (or ‘irregularity’) of neural activity seen in healthy brain function. In this study, we used Multi-scale Rank-Vector Entropy (MRVE) to calculate the dynamic timecourses of signal variability over a range of temporal scales. The correlation of MRVE timecourses was then used to detect functional connections in resting state MEG recordings that were robust over 183 participants and varied with temporal scale. We then compared these MRVE connectivity patterns to those derived using more standard amplitude-amplitude coupling measures, using methods designed to quantify the consistency of these patterns across participants.

Using oscillatory amplitude envelope correlation (AEC), the most consistent connectivity patterns, across the cohort, were seen in the alpha and beta frequency bands. At fine temporal scales (corresponding to ‘scale frequencies’, *f*_*S*_ = 30-150Hz), MRVE correlation detected mostly occipital and parietal connections and these showed high similarity with the networks identified by AEC in the alpha and beta frequency bands. The most consistent connectivity profiles between participants were given by MRVE correlation at *f*_*S*_ = 75Hz and AEC in the beta band.

It was also found that average mid-to fine scale variability within each region (*f*_*S*_ ∼ 10-150Hz) negatively correlated with the region’s overall connectivity strength with other brain areas, as measured by fine scale MRVE correlation (*f*_*S*_ ∼ 30-150Hz) and by alpha and beta band AEC. These findings suggest that local activity at frequencies *f*_*S*_ ≳ 10Hz becomes more regular when a region exhibits high levels of resting state connectivity.

## 1. Introduction

In recent years, MEG has revealed much about the electrophysiological underpinnings of connectivity in the brain. The direct view of neuronal activity provided by MEG and its excellent temporal resolution have allowed the investigation of frequency-specific communication [5] [20] [21] and dynamic changes in connectivity on the millisecond timescale [1] [30].

Alterations in MEG connectivity have also been detected in patient groups [41] [18] [4] [19] [13] [2]. However, to be clinically useful, connectivity research must progress from group-level analysis to the characterisation of individual subjects. To make meaningful comparisons between connectivity profiles of individuals, robust connectivity measures are needed that give consistent results for subjects with the same pathology.

Several recent studies have found that many commonly used techniques for measuring functional connectivity in MEG lack repeatability between healthy subjects, and even show inconsistency over repeated scans of the same subject [45] [8] [25]. [8] found that the method that gave the most consistent connectivity was oscillatory amplitude envelope correlation (AEC), using symmetric orthogonalisation to remove spurious zero-lag correlation between timecourses due to signal leakage [7]. The repeatability of connectivity given by any alternative methods could therefore be compared to AEC to assess the extent to which it can add to our understanding of cortical communication in health and disease.

Many of the most popular techniques for measuring connectivity are based on measuring the synchronisation of oscillatory activity within narrow frequency bands. Another, less studied, aspect of electrical neural activity is the constantly fluctuating activity present in the brain even when it is supposedly ‘at rest’. This variable activity is observed when there is a breakdown of synchrony between neurons, allowing an increase in the information that can be processed within a network [3]. The MEG signals generated by such activity consist of a superposition of many low power signals from smaller neuron populations. This variable activity appears more irregular, or ‘random’, and so is often dismissed as neural ‘noise’, but it is thought to be vital for healthy brain function [11] [17] [36]. It has been shown that the variability of neural activity changes throughout the lifespan [27], and it has been found to be altered in patient groups where activity that is either too regular or too variable is associated with mental disorder [32] [29] [3].

While the physiological role of variability in the brain is not certain, it is possible that it is related to levels of synchronisation between cortical regions, i.e. connectivity. The synchronisation of oscillatory activity, which is highly regular and therefore has low variability, is currently the most promising mechanism for connectivity between brain regions [34] [15] [12] [5] [20] [37]. In contrast, it is thought that local information processing performed within segregated brain regions is associated with signals of high variability that therefore contain higher levels of information [38] [16]. This would lead to the hypothesis that the variability of signals from a cortical region might be related to the levels of connectivity it exhibits with other brain areas.

There is evidence for such a relationship in the literature. One fMRI study found a correlation between the variability of BOLD signals and functional connectivity [44] between spatially separated cortical regions. Age-related connectivity changes have also been shown to covary with the variability of EEG and MEG signals [40] [27] and in an EEG study applying graph theory to functional networks, variability was found to correlate with network node centrality [28].

The variability of neural activity can be quantified using entropy measures, where more disordered and irregular signals have larger entropy, and more regular signals have lower entropy. There are many possible ways of estimating signal entropy [17]. However, one measure that has been shown to be useful in measuring the spatio-temporal variability of MEG signals is Rank-Vector-Entropy (RVE) [33] [3]. RVE is a derivative of Shannon entropy [35] that has a built-in ability to provide a dynamic timecourse of signal entropy, retaining its original temporal resolution. It is also computationally efficient, is calculated from broadband activity timecourses, and is independent of signal amplitude [33]. The relationship between variability and neural synchronisation, and the desirable qualities of RVE, suggest that RVE could be an alternative measure to use in functional connectivity analysis that is not limited to the consideration of oscillatory activity.

RVE, and many other entropy measures, measure signal variability at a single temporal scale. However, it has been shown that neural activity contains recurring patterns that occur across a range of such scales [9]. It is not certain what these correspond to physiologically, however it is thought that activity at coarser scales is associated with long range, distributed information processing, while more local processing is captured at finer scales [40].

To utilise the in-built temporal resolution that is specific to RVE, a multi-scale extension of RVE (MRVE) is proposed [9]. MRVE timecourses at any temporal scale can be calculated from MEG virtual sensor timecourses at any number of required voxels, allowing for a direct comparison with dynamic oscillatory measures. In this paper, MRVE was used used to reconstruct functional connectivity patterns, assess how repeatable these patterns are across a cohort of healthy volunteers and investigate how these patterns vary with temporal scale. We then compared connectivity profiles measured by MRVE correlation with those derived using amplitude envelope correlation (AEC). We also compared the robustness of MRVE correlation, and whether it provides extra information over standard methods, by comparing connectivity patterns derived at multiple entropy time scales with those derived from AEC in multiple frequency bands. Finally, the physiological relevance of variability was investigated by examining the relationships between MRVE, oscillatory amplitude and regional connectivity strength.

## 2. Methods

### 2.1. Data acquisition

Five-minute resting state MEG recordings were acquired from 183 participants (123 female) as part of the ‘100 Brains’ and UK MEG Partnership normative scanning projects. Inclusion criteria ensured all participants were aged 18-65 (mean 24.5±5.4 years), had completed or were undertaking a degree, had normal or corrected-to-normal vision, and had no history of neurological or neuropsychiatric disorders. All procedures were given ethical approval by the Cardiff University School of Psychology Ethics Committee, and all participants gave written informed consent before taking part.

Data were acquired using a whole head 275-channel CTF radial gradiometer system at a 1200 Hz sample rate. An additional 29 reference channels were recorded for noise cancellation purposes and the primary sensors were analysed as synthetic third-order gradiometers [43]. Subjects were seated upright in the magnetically shielded room with their head supported with a chin rest to minimize movement. Participants were asked to rest and fixate their eyes on a central red fixation point, presented on either a CRT monitor or LCD projector. Head localisation was performed at the beginning and end of each scan, using three fiducial markers. Horizontal and vertical electro-oculograms (EOG) were recorded to monitor eye blinks and eye movements.

Participants also underwent a magnetic resonance imaging (MRI) session to acquire a T1-weighted 1mm anatomical scan, using an inversion recovery spoiled gradient echo acquisition (3T, General Electric).

### 2.2. Pre-processing

All data were downsampled to 600Hz and a 1-150Hz bandpass filter applied. Datasets were cut into 2 second epochs, which were each visually inspected and removed if they contained any major artefacts.

Co-registration was performed manually between the MEG and MRI coordinate spaces; the fiducial locations were kept fixed relative to each participant’s nasion, left and right ears and so could then be identified and marked on their MRI scan.

To perform analysis in source space, MEG virtual sensor timecourses were obtained using a scalar LCMV beam-former [42] using FieldTrip [31]. Lead fields were calculated using a localspheres head model for voxels on a 6mm^3^ grid [22]. Covariance matrices were obtained using the broadband pre-processed data filtered between 1-150Hz, as well as for activity within ten narrower frequency bands (1-4, 3-8, 8-13, 13-30, 40-60, 60-80, 80-100, 100-120, 120-140 and 140-160Hz). For all frequency bands, beamformer weights were normalised using the vector norm [20]. The coordinate space for each participant was transformed to the MNI template [14]. The estimated timecourses were then calculated at each voxel for each frequency band, which were then subsequently used to calculate variability and oscillatory amplitude timecourses.

### 2.3 Variability

#### 2.3.1. RVE

The RVE method was first described by [33]. At each time point, a window of *W* points is taken from the signal, each separated by a lag, *ξ*, to avoid oversampling, where *f*_*s*_ represents the sample rate, and *f*_*c*_ is the low-pass frequency applied to the data.

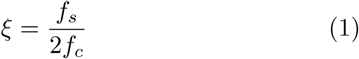

These *W* points are ordered in size, and then converted to the position they originally held in the window. This is the ‘rank-vector’ associated with this time point. The Shannon entropy is calculated at each time point using a state probability distribution derived from the frequency of occurrence of the rank-vectors that occurred previously in the signal [35]. Temporal resolution is introduced using a ‘leaky integrator’, which gives RVE a ‘memory’ of states that is limited in time [33].

#### 2.3.2. MRVE

The calculation of MRVE at each scale is identical to the calculation of RVE, except that each instance of the sliding window is formed from a ‘coarse-grained’ version of the raw signal. For a given scale factor, *S*, at each time point in the signal, *t, W* consecutive, non-overlapping windows of *S* points are taken starting at *t*, where each value is separated by lag *ξ*. Then, the values in the sliding window are found by taking the average of the data points within these windows. This is given by Equation 2, where *x* represents the signal timecourse sampled with lag *ξ*, and ***y***_***t***_ is the window found at time point *t*.

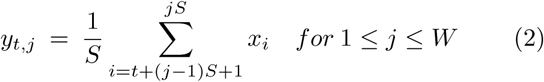

As the rank-vector calculated is dependent on the scale factor used, a separate entropy timecourse is generated for each value of *S* used.

The time scale examined by MRVE is determined by the effective sample frequency of the values in ***y***_***t***_. This ‘scale frequency’, *f*_*S*_, is determined by the scale factor, where a higher value of *S* corresponds to a coarser sampling rate and therefore a lower value of *f*_*S*_ [10]. Equation 3 relates the scale factor to *f*_*S*_ to aid in the interpretation of MRVE and its relationship with oscillatory measures.

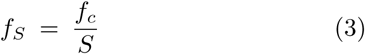

### 2.4. Functional connectivity

90 nodes were selected by taking one voxel timecourse to represent each region of the AAL atlas [39]. The selection was performed for each participant and for each frequency band, by identifying the virtual sensor time course, within each AAL region, that had the highest temporal standard deviation. To avoid the detection of spurious connections due to signal leakage, the zero-lag correlation between all 90 AAL timecourses was removed by symmetric orthogonalisation [7]. This resulted in 90 orthogonal timecourses for each participant and frequency range, which were then used to calculate MRVE and oscillatory amplitude timecourses.

MRVE was calculated from the broadband, 1-150Hz virtual sensor timecourses, using a window length of *W* = 5 and a decay time constant of *τ* = 0.07*s*. Timecourses were calculated for 25 scale factors between *S* = 1 − 150, with corresponding scale frequencies ranging from *f*_*S*_ = 1 − 150Hz. Oscillatory amplitude envelopes were found by applying the Hilbert transform to the timecourses obtained for each of the aforementioned narrow frequency bands. Functional connectivity was then measured as described by [23]. The MRVE and Hilbert envelope timecourses were de-spiked to remove artefactual temporal transients using a median filter, and downsampled to 1 Hz. The first 50 samples were then trimmed to remove the MRVE ‘warmup’ period while the histogram populates, and a window of samples at the end was removed, the length of which was defined by the length in time of the longest sliding window used in the MRVE calculation, corresponding to the largest scale factor.

Functional connectivity matrices were calculated separately for MRVE at each scale, and for oscillatory amplitude within each narrow frequency band by correlating each of the 90 timecourses from each participant with all others. The correlation values were then normalised by converting them to Z-scores using the Fisher transform. These were variance-normalised to correct for the effects of the varying timecourse lengths between participants, due to the removal of data epochs containing artefacts [23]. Significant connections were determined by first ranking connections in order of strength for each participant, where the strongest connection was given the value 1 and the weakest given value 0. ‘Valid connections’ were taken as those with a mean rank above a threshold of 0.8, indicating that these connections are consistently among the strongest across participants. This threshold is arbitrary, however it has been shown previously to be a suitable threshold for detecting robust resting state network connections using AEC [23].

### 2.5. Software

Data analysis was performed in MATLAB, using Fieldtrip functions and custom built MATLAB scripts [31]. Connections were visualised on a template brain using the SourceMesh MATLAB toolbox, and voxel-wise correlation colourmaps were created in mri3dX.

## 3. Results

### 3.1. Consistency of functional connectivity across participants

First, MRVE correlation was used to measure functional connectivity and we assessed which of these connections were consistently among the strongest across subjects. Figure 1 shows the location and number of the valid connections found for scale frequencies, *f*_*S*_ = 1-150Hz. At higher scale frequencies, i.e. at finer temporal scales, most connections are found in occipital and parietal regions. As shown in Figure 2, the maximum number of connections was found at *f*_*S*_ = 75Hz. However, there is a second peak in the number of valid connections found at *f*_*S*_ = 10Hz, where more frontal connections are seen. Cumulatively across all scales, valid connections were detected between 254 different pairs of nodes.

**Figure 1:**
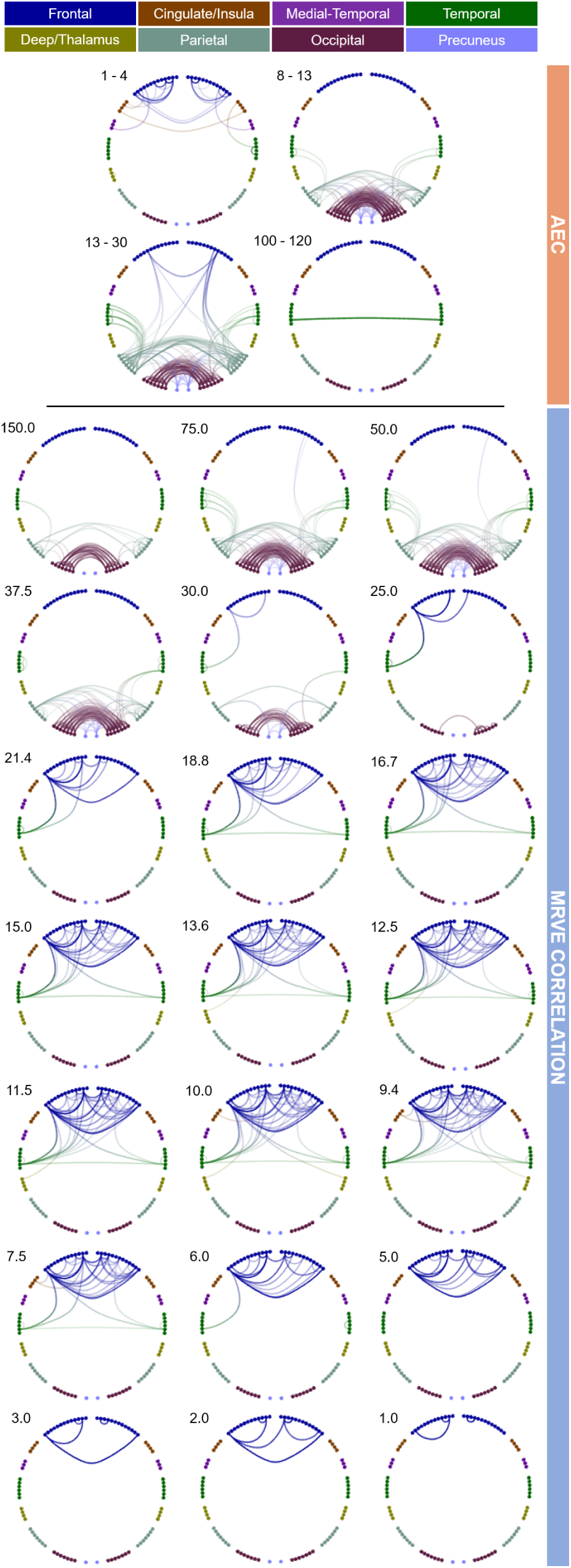
Valid connections (mean rank > 0.8) found using AEC correlation for four frequency bands (above) and MRVE correlation for a range of time scales (below). Each point represents an AAL region and each line represents a connection. The midpoint of the frequency band (for AEC) or scale frequency (for MRVE correlation) is indicated in the top left corner of each plot, in Hz. The key at the top indicates the colour of the connections that originate in each brain region. No valid connections were found using AEC in the 3-8, 40-60, 60-80, 80-100 or 140-160Hz frequency bands.

**Figure 2:**
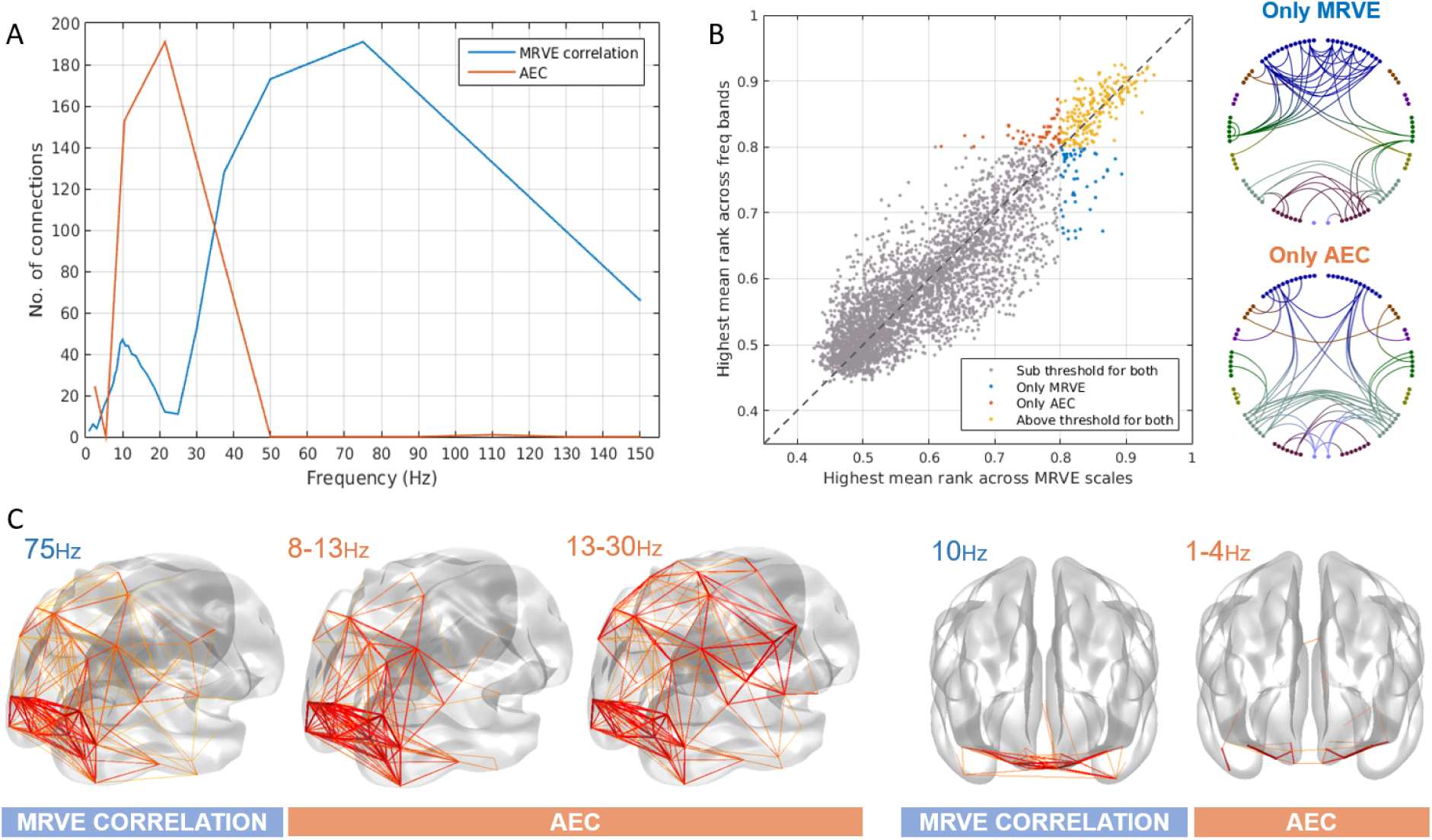
A) The number of valid connections found for each scale frequency using MRVE correlation and each frequency band using AEC. B) The highest mean rank of each connection across all frequency bands vs. all scale frequencies, where the colour indicates whether the connection is detected by both MRVE correlation and AEC, detected by neither, or detected by only one of the methods. Circle plots show the connections that are only detected as valid by either MRVE correlation or AEC. C) Valid connections plotted on a template brain for MRVE scale frequencies *f*_*S*_ = 75Hz and 10Hz and AEC frequency bands 1-4Hz, 8-13Hz and 13-30Hz.

The valid connections found using AEC are also shown in Figure 1. Valid connections were found within four frequency bands. The most valid connections were seen in the beta band, giving the same number as for *f*_*S*_ = 75Hz using MRVE correlation. Across all frequency bands, valid connections were detected between 248 different node pairs.

We then investigated whether MRVE correlation could provide additional information about functional connectivity beyond that provided by AEC analysis. Firstly, it was seen whether each method could detect unique connections that were not deemed valid by the alternate method. To determine this for each connection, its mean rank, averaged across all subjects, was calculated for all scale frequencies for MRVE and for all frequency bands using AEC. For each connection, we then found the highest mean rank for any of the MRVE scales and the highest mean rank for any of the AEC frequency bands. Those with a highest mean rank above the threshold of 0.8 for either method were taken as detectable by the corresponding connectivity measure. Figure 2 shows the highest mean rank values for each connection plotted against each other. Those connections that are ‘unique’ to each method are shown plotted between the AAL nodes. 200 connections are visible for both MRVE correlation and AEC across all scales and frequency ranges, leaving 54 connections (21%) that can only be seen using MRVE correlation, and 49 (19%) that can only be seen using AEC.

#### 3.1.1. Robustness of connectivity measures to sample size

The robustness of each connectivity measure to the participant sample size was determined using bootstrapping. Sub-samples of a range of sizes were taken from the participant cohort by simple random sampling with replacement. The number of valid connections was found for each sub-group taken, over 1000 tests per sub-group size, *N*. It can be seen in Figure 3A that the average number of connections found was less stable when using fewer participants in the analysis for both MRVE correlation and AEC. The average number approximates to the number of connections detected using the whole cohort (as shown in Figure 2a) when using *N* ≳ 60. However, for both MRVE correlation and AEC, the variance in the number of valid connections detected was found to be larger when fewer participants were included. For *N* ≲ 60, a smaller sample was also associated with more connections detected on average.

**Figure 3:**
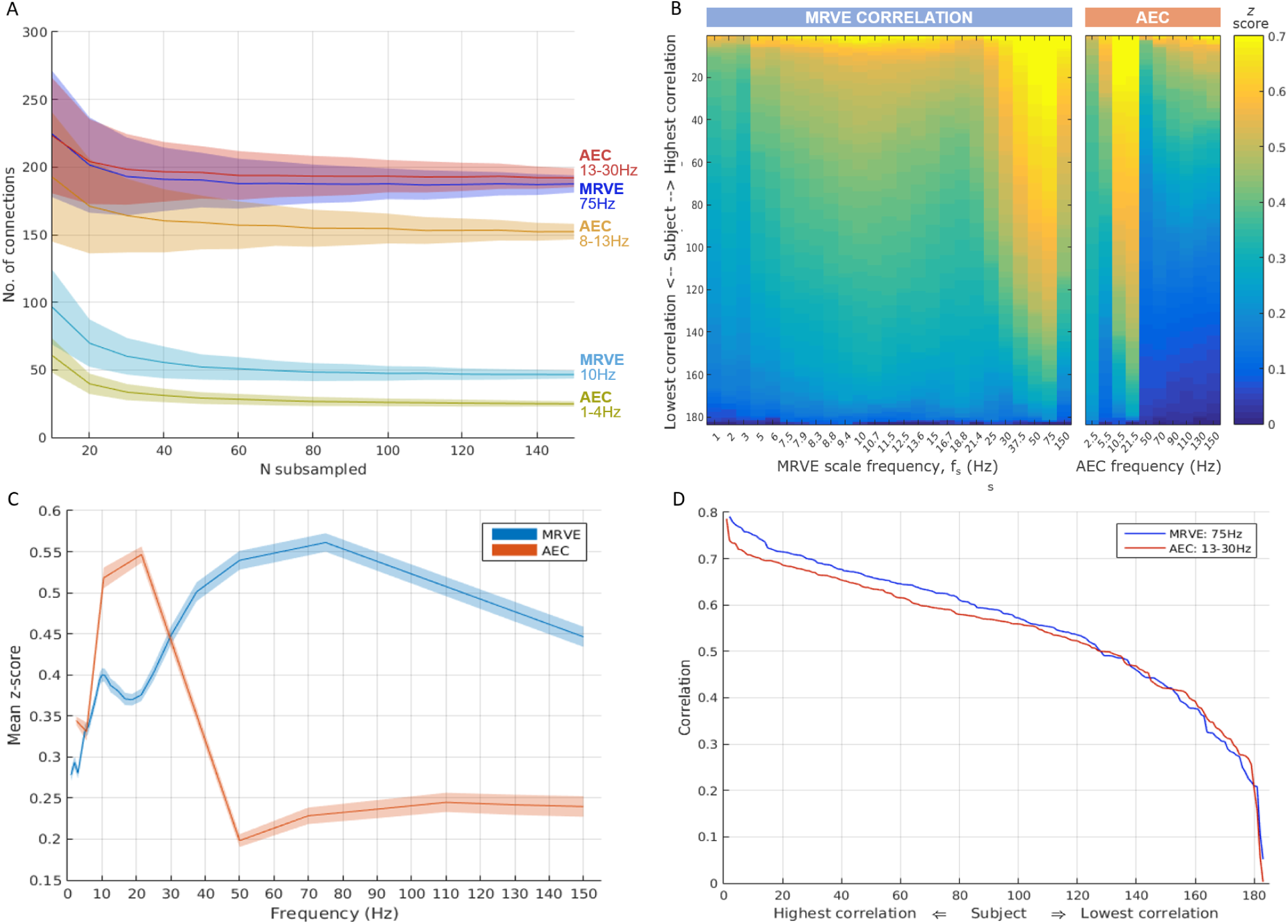
Analysis of the robustness and inter-participant consistency of the connectivity measures given by AEC and MRVE correlation. A) Robustness to reducing the number of subjects included in the analysis as measured by bootstrapping. Plot shows the mean number of valid connections detected over 1000 sub-samples of size *N*, randomly sampled with replacement. Error bands show the standard deviation. B) Consistency across subjects found by correlating the vectorised *z* score connectivity matrices of individual subjects with the average connectivity pattern across all subjects. Colour plots show the resultant *pattern-correlation* coefficients for each subject, sorted by correlation strength, for each MRVE scale frequency and AEC frequency band. AEC bands are represented by the frequency at the midpoint between the limits of the frequency range. C) These *pattern-correlation* coefficients over subjects were transformed to *z* scores and averaged for each scale frequency and frequency band. Error bands show the standard deviation over 1000 sub-samples of *N* = 90. D) Sorted *pattern-correlation* coefficients, calculated as in B) for MRVE scale frequency *f*_*S*_ = 75Hz and beta band AEC.

#### 3.1.2. Consistency of connectivity patterns across participants

The consistency of the connectivity profiles between individuals was then investigated. The average connectivity profile for each frequency band and scale frequency was taken by vectorising the mean *z* score connectivity matrix. This profile was then correlated with the equivalent vector of *z* scores obtained for each participant individually. For very robust networks that are highly reproducible across subjects, this method will give consistently high *pattern-correlation* with the average connectivity profile. However, the distribution of correlation coefficients will be, on average, lower for a network that shows high variability across participants. Each *pattern-correlation* coefficient is represented in the colour plot shown in figure 3B. For each scale factor and frequency band, these have been sorted in descending order of participants. Consistent high correlation with the average connectivity patterns, representing high cross-subject repeatability, can be seen for MRVE correlation at scale frequencies 50 and 75Hz, and for alpha and beta band AEC.

To further quantify the consistency of each connectivity measure across subjects, the mean correlation with the average connectivity profile was found for each scale frequency and frequency band (i.e. the average was taken from each column on the colour plot). These average *pattern-correlation* values are shown in figure 3C, with errorbands generated by bootstrapping, using 1000 sub-samples of group size *N* = 90. The highest mean *pattern-correlation* was found for MRVE correlation, *f*_*S*_ =75Hz (*r* = 0.5089 ± 0.0004), followed by beta band AEC (*r* = 0.4980 ± 0.0003), suggesting that these two connectivity profiles were the most reproducible across subjects.

The *pattern-correlation* values (as shown in Figure 3b) were then compared for MRVE correlation at *f*_*S*_ = 75Hz and beta band AEC. The sorted *pattern-correlation* values for these frequencies are shown in Figure 3D. It appears that MRVE correlation at *f*_*S*_ = 75Hz gives individual profiles that are slightly more similar to the average connectivity profile than beta band AEC. The *pattern-correlation* values were then compared in a permutation test, where the group assignment was randomised between 75Hz MRVE and beta band AEC over 10,000 permutations. However, it was found that there was no significant difference between the *pattern-correlation* values (p=0.344) for each connectivity measure.

#### 3.1.3 Predicting MRVE connectivity from AEC connectivity

The amount of variance in the MRVE correlation that could be explained by AEC was then calculated using a multiple regression model. The fraction of the variance in the MRVE connectivity that could be explained by AEC was then calculated by vectorising the connectivity matrices and using the model in equation 4, where *i* represents each frequency band, *N*_*f*_ is the number of frequency bands used in the model and *x*_*i*_ represent the regression coefficients.

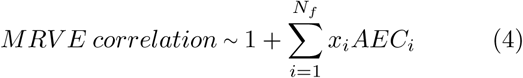

The adjusted R^2^ value found for each scale factor is shown in Figure 4. The adjusted R^2^ value was used to determine which combination of frequency bands would best explain the MRVE connectivity, as the highest adjusted R^2^ values are obtained when the model only includes predictor variables which add explained variance beyond that which would be expected by chance. However, it was found that the highest adjusted R^2^ values for each scale frequency were achieved when the AEC connectivity vectors from all frequency bands were incorporated in the model, except for *f*_*S*_ = 7.9Hz when the alpha band was excluded, and for *f*_*S*_ = 150Hz when the 80-100Hz band was excluded.

**Figure 4:**
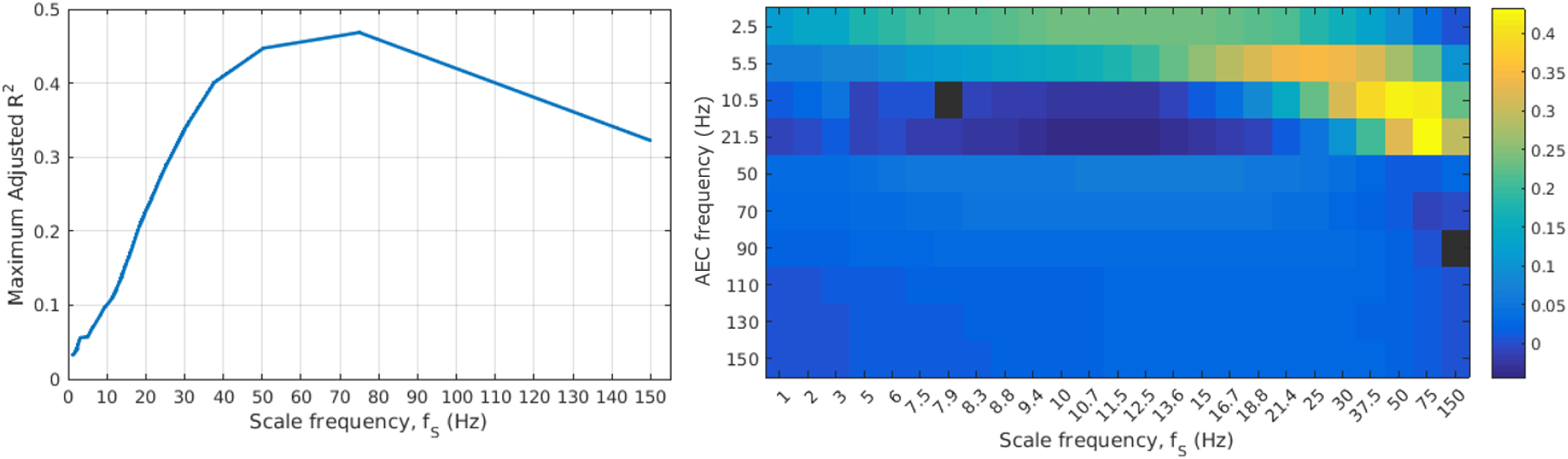
(left) The maximum adjusted R^2^ obtained across all possible multiple linear regression models for each scale frequency. (right) Colour plot showing regression coefficients, where each column represents the coefficients obtained using MRVE correlation at the given scale frequency as the response variable. AEC bands are represented by the frequency at the midpoint between the limits of the frequency range. Black indicates that the corresponding AEC frequency was not included as a predictor variable in the optimal regression model (maximising adjusted R^2^).

### 3.2. Temporal correlation between MRVE and oscillatory amplitude envelopes

The relationship between entropy and oscillatory amplitude was then investigated. At each voxel in the brain, the temporal correlation between MRVE timecourses and oscillatory amplitude envelopes was found across scale frequencies and frequency bands. Average *z*-scores are shown on a template brain in Figure 5.

**Figure 5:**
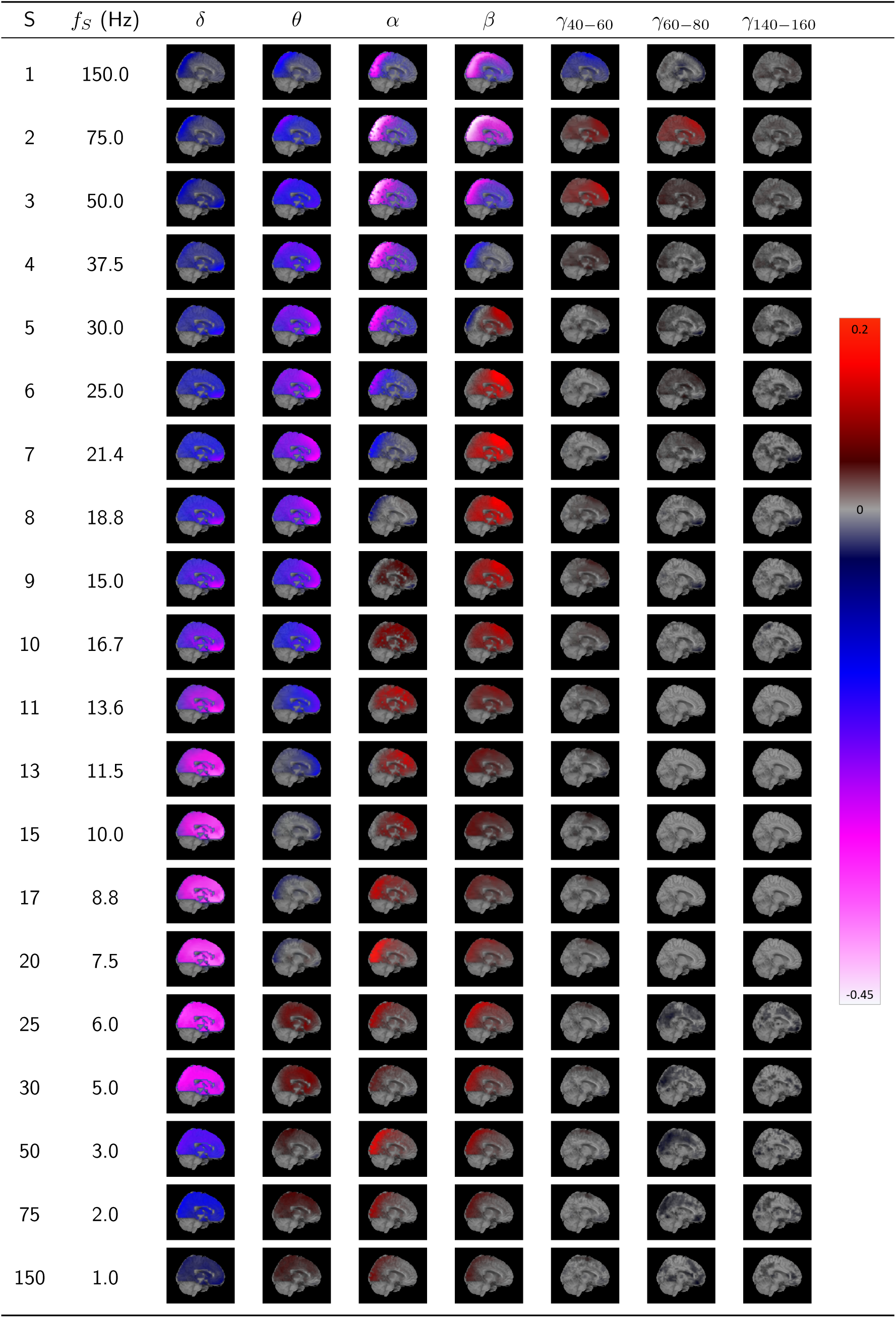
The temporal correlation between MRVE timecourses and oscillatory amplitude envelopes for scale frequencies *f*_*S*_ = 1-150Hz and frequency bands 1-4Hz (*δ*), 3-8Hz (*θ*), 8-13Hz (*α*), 13-30Hz (*β*), 40-60Hz (*γ*_40−60_), 60-80Hz (*γ*_60−80_) and 140-160Hz (*γ*_140−160_). The temporal correlation coefficient was found at each voxel for each participant and transformed to a z-score by applying the Fisher transformation. The 95% confidence interval was found for the *z*-scores calculated across all participants for each voxel. Average Pearson correlation values were found at each voxel where *z* = 0 lay outside of this confidence interval and displayed on a template brain as indicated by the colour bar. See supplementary material for whole brain correlation images for all scales and frequency bands.

The relationship is shown to be dependent on the MRVE scale frequency and oscillatory frequency band. However, the direction is generally consistent across the brain for each combination. At high scale frequencies (*f*_*S*_ = 50-150Hz), MRVE shows a strong negative correlation with power in the alpha and beta frequency bands, where the strongest relationship is seen between MRVE at *f*_*S*_ = 75Hz and beta band amplitude in the occipital and parietal regions. At *f*_*S*_ = 50-75Hz, a weak positive correlation with gamma band amplitude is also observed, which is strongest in frontal and temporal regions. At mid to lower scale frequencies (*f*_*S*_ = 1-25Hz), MRVE shows a negative correlation with delta band amplitude but a positive correlation with power in the alpha and beta bands. However, the areas in which the strongest positive correlation is observed varies with scale frequency and differs between the two frequency bands. The strongest positive correlation was observed between MRVE at *f*_*S*_ = 21.4Hz and beta band amplitude in frontal and temporal regions. However, positive correlation was also observed in occipital and parietal regions between alpha and beta band amplitude and MRVE for *f*_*S*_ = 1-8.8Hz.

### 3.3 The relationship between MRVE magnitude, oscillatory amplitude and connectivity strength

The overall connectivity ‘strength’ was then estimated for each AAL region. This was done for each node by finding the sum of the correlation coefficients indicating the connectivity between that node and all other nodes, for each AEC frequency band and each MRVE correlation scale frequency. This gave one connectivity strength value for each AAL region for each participant, for each frequency band and scale frequency used. The average entropy value within each AAL region was then found at each scale frequency for each participant, by taking the average value of the MRVE timecourses from the node voxel used in the connectivity analysis. Figure 6 shows the correlation between a vector containing all average entropy values across participants and the corresponding connectivity strength values. The correlation between connectivity strength and average oscillatory amplitude was also found, taken as the mean value of the hilbert envelope calculated for activity in each frequency band.

**Figure 6:**
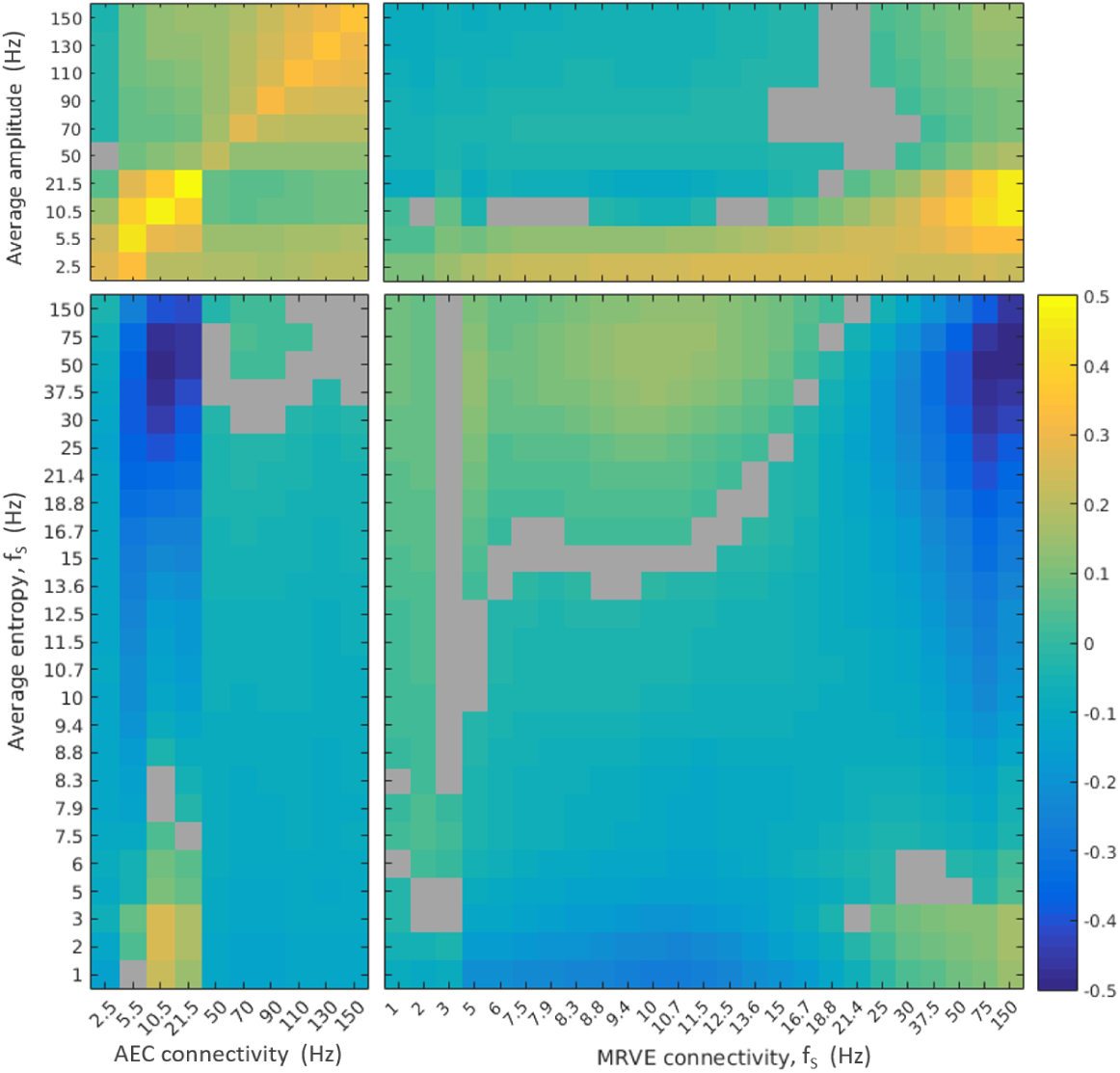
The correlation between average oscillatory amplitude/entropy and the overall connectivity strength at each voxel as measured by AEC and MRVE correlation. AEC bands are represented by the frequency at the midpoint between the limits of the frequency range. Warm colours indicate positive correlation whereas cooler colours show negative correlation and grey indicates a non-significant relationship.

At high scale frequencies, it was generally found that average variability negatively correlates with connectivity strength. The strongest relationship with MRVE correlation was found between average entropy at *f*_*S*_ = 75Hz and connectivity strength at *f*_*S*_ = 150Hz (*r* = −0.62, *p* ≪ 0.001), whereas the strongest relationship with AEC was found between average entropy at *f*_*S*_ = 50Hz and alpha band connectivity strength (*r* = −0.50, *p* ≪ 0.001). However, a weaker positive correlation was found between average entropy at fine time scales and connectivity at coarser scales, where the strongest correlation was found between average entropy at *f*_*S*_ = 75Hz and connectivity strength at *f*_*S*_ = 10Hz (*r* = 0.18, *p* ≪ 0.001). A positive correlation is also seen between average entropy at very low scale frequencies (*f*_*S*_ = 1-3Hz) and AEC in the alpha and beta bands, as well as with MRVE correlation at the highest scale frequencies. This is strongest between average entropy at *f*_*S*_ = 3Hz and alpha band AEC (*r* = 0.25, *p* ≪ 0.001), and between average entropy at *f*_*S*_ = 2Hz and MRVE correlation at *f*_*S*_ = 150Hz (*r* = 0.20, *p* ≪ 0.001).

In contrast, there was generally a positive relationship between average oscillatory amplitude and connectivity strength. As shown in the top left of Figure 6, for AEC connectivity the strongest correlations were found when relating amplitude and connectivity strength within the same frequency band, where the strongest relationship was found for the beta band (*r* = 0.49, *p* ≪ 0.001). Average amplitude also generally showed a positive correlation with connectivity strength as measured by MRVE correlation at fine time scales, where the strongest relationship was found between alpha band amplitude and connectivity strength for *f*_*S*_ = 150Hz (*r* = 0.46, *p* ≪ 0.001).

## 4. Discussion

The correlation of neural variability as measured by MRVE was used here to detect robust functional connections from MEG recordings, suggesting that this is a viable method for the analysis of resting state connectivity. The existence of robust connections that can only be detected by MRVE correlation also suggests that this method can provide complementary information to that provided by AEC.

By introducing the multi-scale element to the RVE method, it was possible to observe network connections that were present at different temporal scales. The number of valid connections detected and the brain areas they originated from varied with each scale frequency, although it was found that two general patterns of connectivity emerged.

At finer temporal scales (*f*_*S*_ = 30-150Hz), the networks revealed are dominated by occipital and parietal connections, with some fronto-parietal and temporo-parietal connections. Connectivity in these regions during the resting state has been well established in the literature, in both fMRI [24] and MEG studies, where connections in these areas have been found in the alpha and beta frequency bands using oscillation-based connectivity measures [5] [20].

The relationship between MRVE at fine time scales and oscillatory amplitude in the alpha and beta frequency bands was a recurring feature throughout the analysis here. It was shown that the connectivity profiles revealed by fine-scale MRVE correlation and AEC in the alpha and beta bands showed high levels of similarity; the AEC within the alpha and beta frequency ranges made large contributions to the explained variance in the MRVE correlation at *f*_*S*_ = 75Hz, the scale at which most connections were detected. It was also found that fine-scale variability time-courses exhibited a strong negative correlation with the alpha and beta band amplitude envelopes, and that connectivity strength negatively correlates with average MRVE at this frequency while positively correlating with alpha and beta band amplitude. These findings imply that high levels of alpha and beta band AEC are associated with more regular activity at scale frequencies *f*_*S*_ = 30-150Hz.

It could be that the decrease in variability represents a reduction in information processing performed locally within areas showing high levels of inter-regional connectivity. Entropy is maximised when there is the least integration between brain regions, while increased connectivity introduces statistical dependencies from activity in other brain areas and so decreases the ‘randomness’ exhibited by a region [38].

These results are also potentially consistent with a computational model that recently showed that correlated amplitude envelope fluctuations in the alpha and beta bands are driven by time-delayed coupling between oscillators in the gamma band [6]. It was found that transient synchronisation between these oscillators led to correlated amplitude fluctuations at a reduced collective frequency. Future work could investigate whether the correlation between entropy timecourses at high scale frequencies is driven by the degree of synchronisation between oscillators at the same natural frequencies.

At coarser temporal scales, a second network pattern emerged consisting of mostly frontal and temporal connections that most closely resembled the AEC network found within the delta band. This similarity was again supported by the regression analysis, where the delta band AEC explained the largest fraction of variance in the MRVE correlation for scale frequencies *f*_*S*_ = 1 − 13.6Hz. However, the overall fraction of the variance that can be explained at these coarser time scales is relatively small, suggesting that MRVE correlation provides more novel information about connectivity at these scales beyond that which can be observed by AEC. Although, as shown in Figure 2c, the connections detected at lower scale frequencies originated at the very front of the brain. While all datasets were visually cleaned and each set of virtual sensor timecourses were orthogonalised, it is possible that contamination by signals from eye movements could be causing spurious connections in these regions. Future work could repeat the analysis outlined here using data that has been cleaned of eye movement artefacts, for example using ICA.

It is interesting to note that MRVE correlation, for a given scale frequency, does not provide the same information about functional connectivity as AEC for an overlapping frequency band. For example, while the frequency band that shows the most connections using AEC ranges from 13-30Hz, scale frequencies in this range are associated with a trough in the number of connections when using MRVE correlation. In fact, Figure 4 shows that for each frequency band, AEC explains a low percentage of the variance in the MRVE correlation at scale frequencies in the same range. This suggests that in regions showing high connectivity strength, the amplitude and variability of activity of a particular frequency are not related.

However, MRVE was shown to have a complex relationship with oscillatory amplitude. In general, a positive correlation was found between oscillatory amplitude and entropy timecourses calculated within the same frequency range, whereas a negative correlation is seen when the MRVE scale frequency is approximately higher than the lowpass frequency of the oscillatory frequency band. At the finest time scales, this is seen as a biphasic relationship where MRVE shows negative correlation with low frequency amplitude but positive correlation with gamma band amplitude. This has been found previously in a study using RVE at a single time scale, when applied to task data (*f*_*S*_ = 150Hz) [3]. Here, the relationship was replicated in resting state data and was found to be consistent in direction across the brain. However, by considering multiple time scales using MRVE, it was found that the correlation between entropy and amplitude envelopes varies with the entropy scale frequency.

While the direction of each relationship was found to be generally consistent across the brain, the strength of the relationships were often found to vary spatially. For a number of combinations, the correlation was found to be strongest either in occipital and parietal regions, where most functional connections were detected, or in frontal and temporal regions. For example, the negative correlation between beta band amplitude and MRVE at fine scales is strongest in more posterior regions. In contrast, the positive correlation observed for *f*_*S*_ values within the beta frequency range is strongest in anterior regions. This could imply that regional connectivity strength moderates the relationship between the variability and oscillatory amplitude of neural activity within that region. Future work could look at whether the same phenomenon is observed during a task, during which different regions would show higher connectivity strength.

Connectivity strength was generally found to positively correlate with oscillatory amplitude, in agreement with previous research [37], but was found to negatively correlate with variability. This is consistent with the prevailing theory that oscillatory activity (which is highly regular) facilitates synchronisation between cortical regions [34] [15] [6]. The relationship between connectivity and oscillatory amplitude is often confounded by the fact that an increase in amplitude is associated with an increase in SNR. However, it is unlikely that this would be causing the observed relationship with variability. If the low measured entropy was driven by increases in underlying signal strength, we would expect to detect connections in the gamma band using AEC that match those found by MRVE correlation for scale frequencies in the same range, whereas in reality very few connections are seen using AEC at these high frequencies.

While a weaker positive correlation was observed between connectivity at coarse scales and average entropy at fine scales, it is uncertain whether connections at these scales were spurious due to eye movement artefacts. Therefore it is uncertain whether this relationship would be replicated if these were removed.

A limitation of this study is that the performance of MRVE correlation was only compared to AEC. AEC was chosen for comparison as it has been shown to give the most consistent results across participants [8]. This method was therefore the appropriate benchmark to use in a comparison of the number of robust connections detected by each connectivity measure. However, it could also be interesting to look at how MRVE correlation relates to connectivity measured by techniques that are centred around phase relationships. It has been suggested that the reduction of signal variability facilitates phase relationships to occur between brain regions [26] so it could be investigated whether there is similarity between the connections each method can detect and how this would differ from the relationship between MRVE correlation and AEC.

Another constraint on this analysis was the limit on the resolution of scale frequencies that could be used to generate MRVE timecourses. By increasing the sample rate of the data acquisition, it would be possible to obtain MRVE timecourses at finer temporal scales, and at more frequencies within the frequency range investigated here. For example, with a lowpass frequency of 300Hz, MRVE timecourses could be calculated for all frequencies considered here, as well as for *f*_*S*_ = 300Hz, 100Hz, 60Hz etc. Future work could therefore choose the sample rate and scale factors used in order to target specific frequencies of interest.

While it is interesting that MRVE correlation has shown promise as a measure of functional connectivity, the true test of its usefulness will be its performance in patient groups. Neural variability measures, such as Multi-scale Entropy (MSE) [9], and AEC connectivity have both been shown to be able to distinguish patient groups from controls. Future work will investigate whether MRVE correlation can provide understanding about connectivity changes associated with disease, in comparison to conventional measures based on the oscillatory components of brain function.

## 5. Acknowledgements

This research was supported by CUBRIC and the School of Psychology at Cardiff University, the UK MEG Partnership Grant (MRC/EPSRC, MR/K005464/1) and EPSRC studentship funding to Godfrey.

