## Supplemental to Figure 5 for "Measuring robust functional connectivity from resting-state MEG using amplitude and entropy correlation across frequency-bands and temporal scales"



**Abstract**

Supplementary figures to complement Figure 6 in the main article. The relationship between MRVE time-courses and oscillatory amplitude is shown here across the whole brain, including all frequency bands and MRVE scale frequencies used in the connectivity analysis.


| S | $f_S$ (Hz) | $\delta$ | $\theta$ | $\alpha$ | $\beta$ | $\gamma_{40-60}$ |
|---|---|---|---|---|---|---|
| 1 | 150.0 | | | | | |
| 2 | 75.0 | | | | | |
| 3 | 50.0 | | | | | |
| 4 | 37.5 | | | | | |
| 5 | 30.0 | | | | | |
| 6 | 25.0 | | | | | |
| 7 | 21.4 | | | | | |
| 8 | 18.8 | | | | | |
| 9 | 16.7 | | | | | |
| 10 | 15.0 | | | | | |

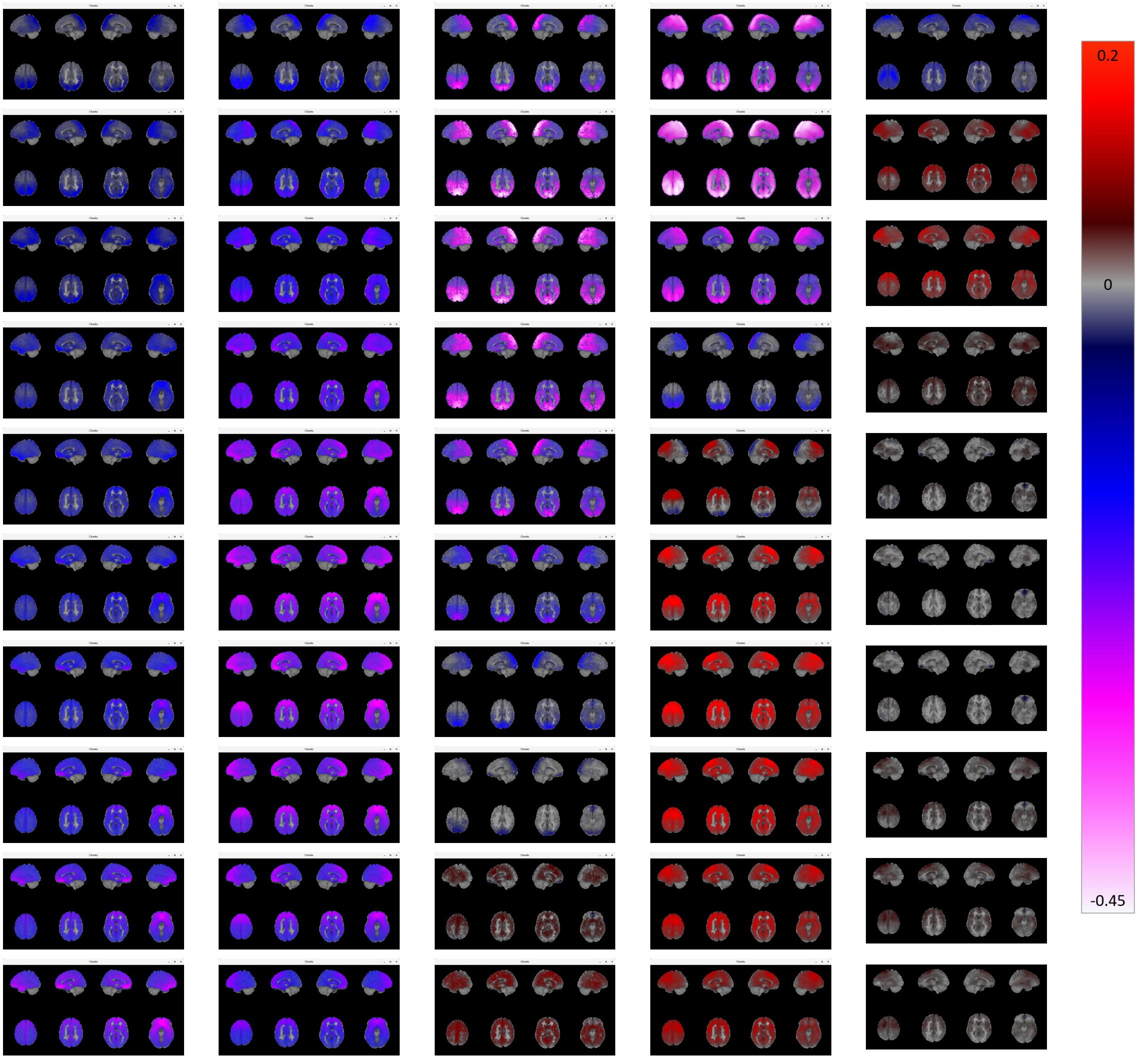


*Preprint submitted to Elsevier*

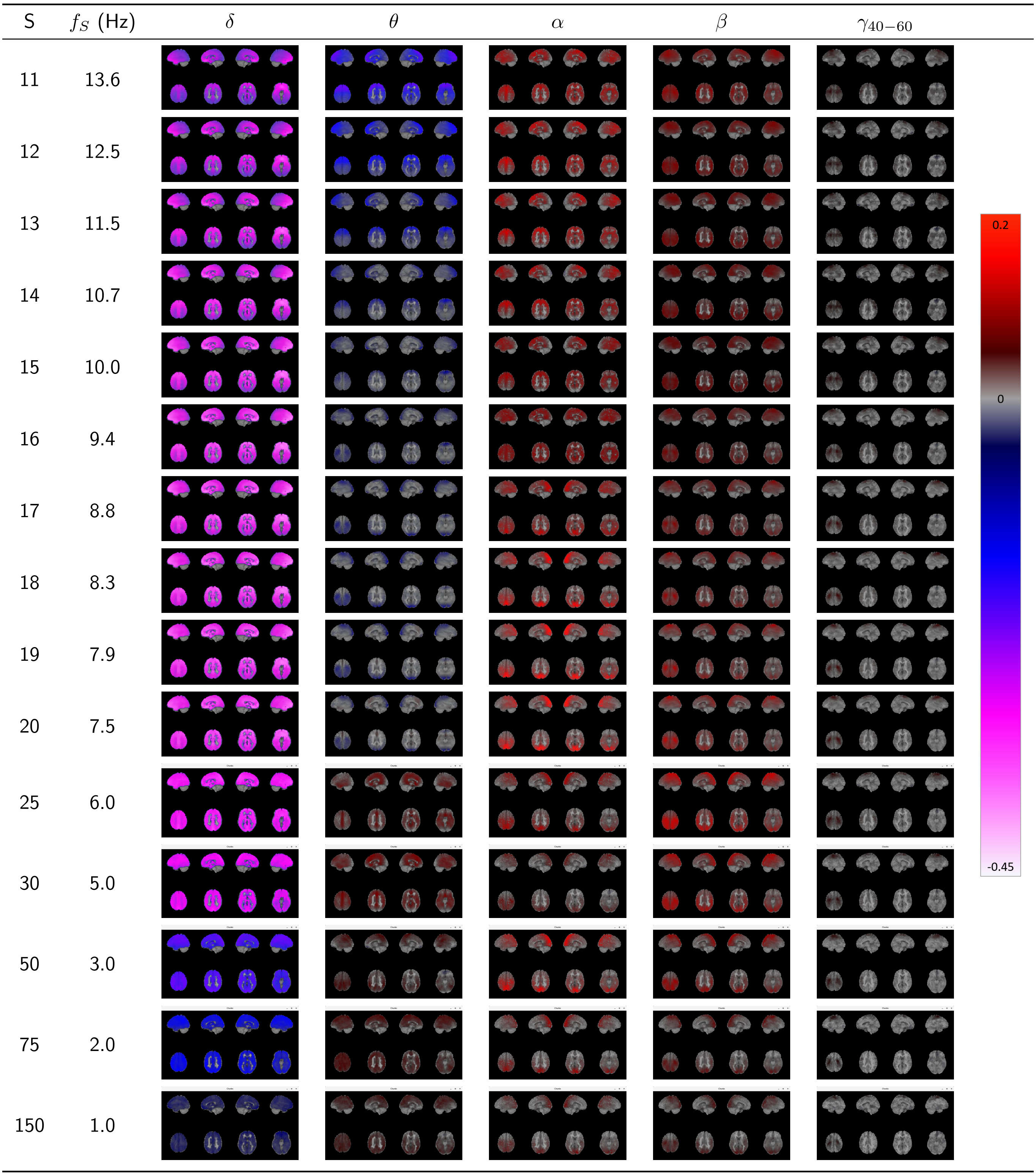


Figure 1: The whole-brain correlation between MRVE time-courses and oscillatory amplitude envelopes for scale frequencies $f_S$ = 1-150Hz and frequency bands 1-4Hz ($\delta$), 3-8Hz ($\theta$), 8-13Hz ($\alpha$), 13-30Hz ($\beta$) and 40-60Hz ($\gamma_{40-60}$). Correlation was found at each voxel for each participant and transformed to a z-score by applying the Fisher transformation. The 95% confidence interval was found for the $z$-scores calculated across all participants for each voxel. Average Pearson correlation values were found at each voxel where $z = 0$ lay outside of this confidence interval and displayed on a template brain as indicated by the colour bar.

| S | $f_S$ (Hz) | $\gamma_{60-80}$ | $\gamma_{80-100}$ | $\gamma_{100-120}$ | $\gamma_{120-140}$ | $\gamma_{140-160}$ |
|---|---|---|---|---|---|---|
| 1 | 150.0 | | | | | |
| 2 | 75.0 | | | | | |
| 3 | 50.0 | | | | | |
| 4 | 37.5 | | | | | |
| 5 | 30.0 | | | | | |
| 6 | 25.0 | | | | | |
| 7 | 21.4 | | | | | |
| 8 | 18.8 | | | | | |
| 9 | 16.7 | | | | | |
| 10 | 15.0 | | | | | |
| 11 | 13.6 | | | | | |
| 12 | 12.5 | | | | | |
| 13 | 11.5 | | | | | |
| 14 | 10.7 | | | | | |
| 15 | 10.0 | | | | | |

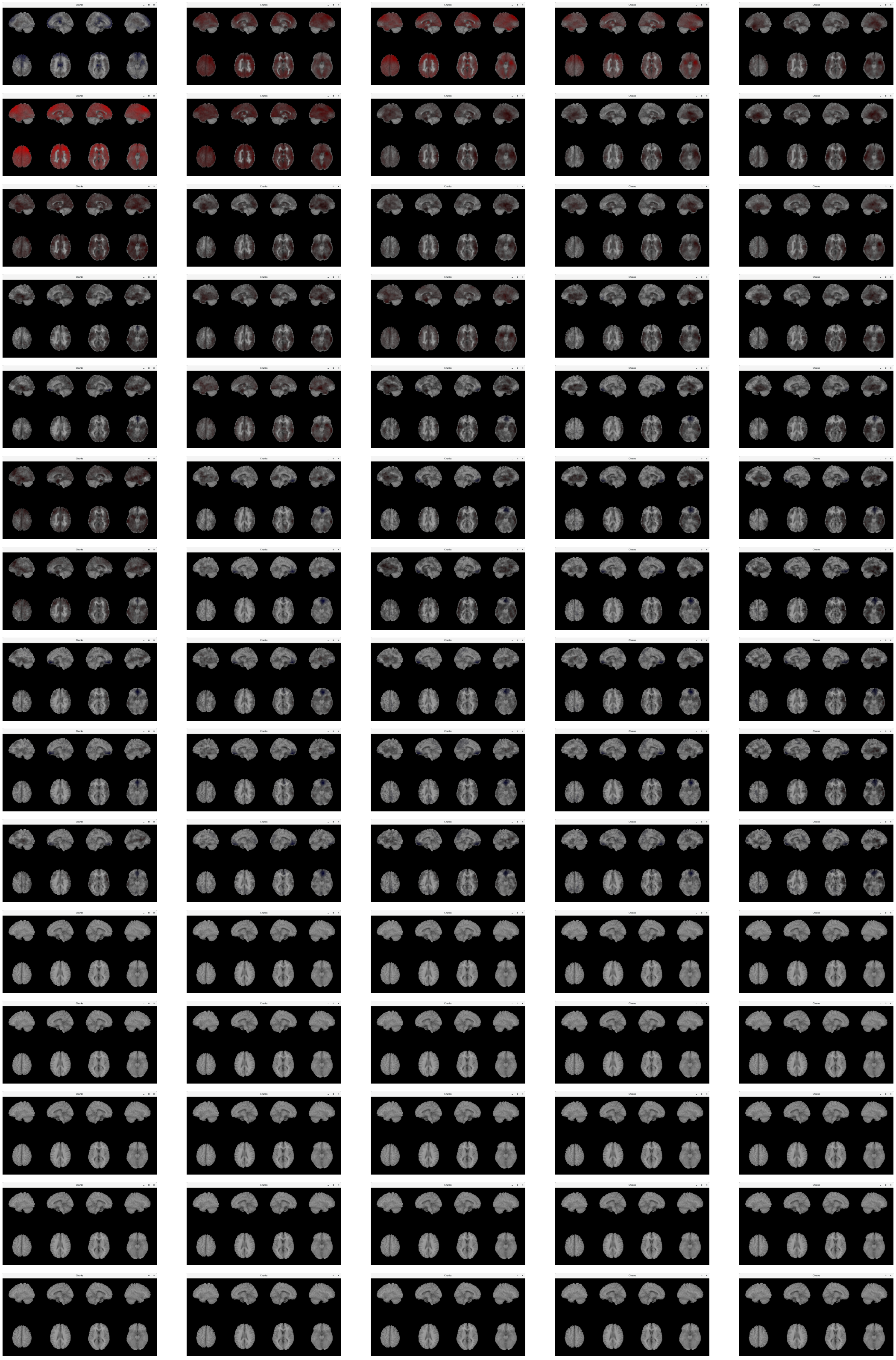

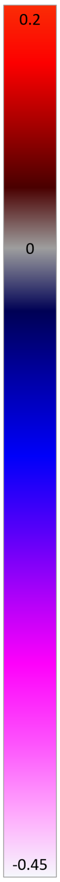

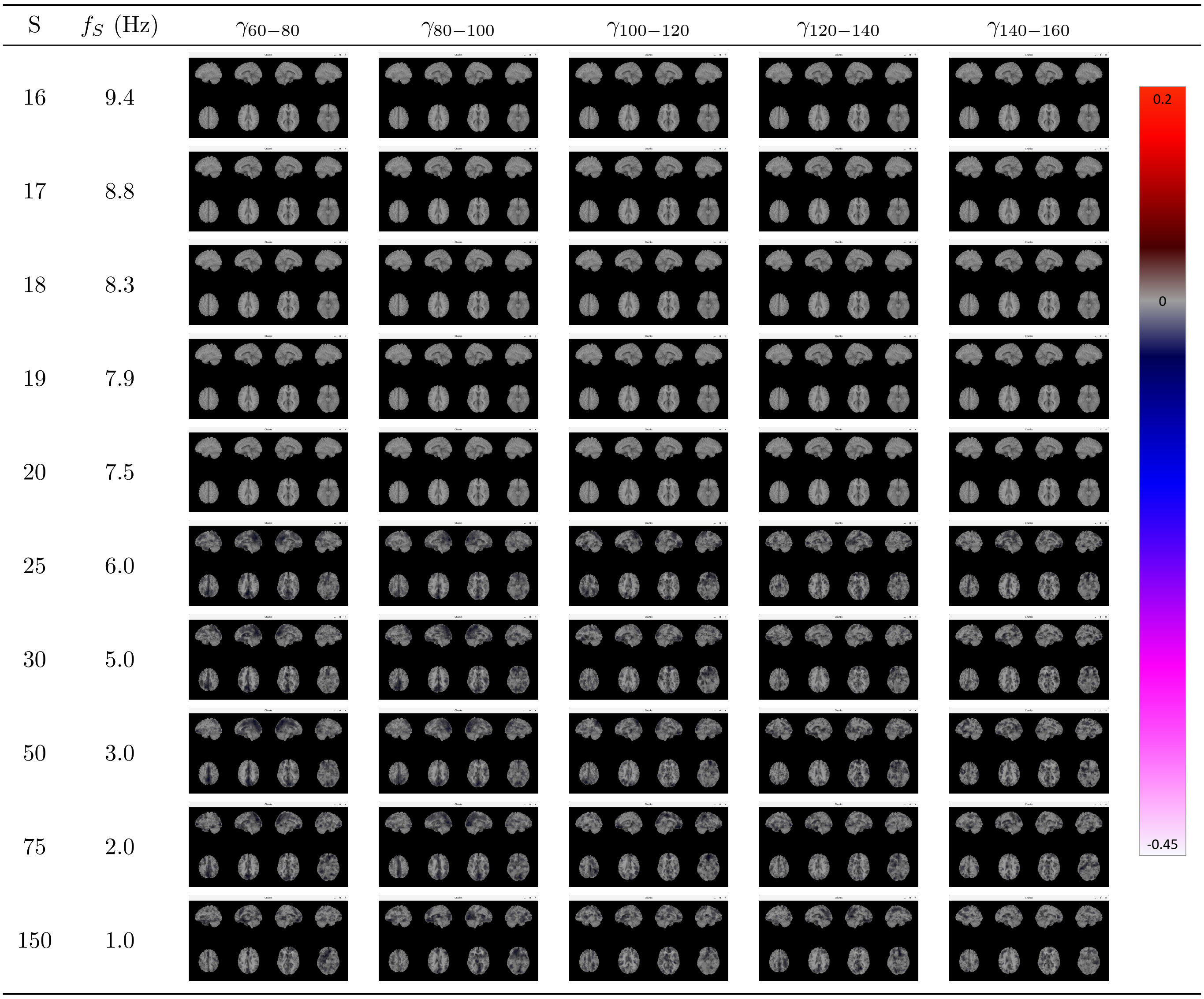


Figure 2: The whole-brain correlation between MRVE time-courses and oscillatory amplitude envelopes, for scale frequencies $f_S$ = 1-150Hz and frequency bands 60-80Hz ($\gamma_{60-80}$), 80-100Hz ($\gamma_{80-100}$), 100-120Hz ($\gamma_{100-120}$), 120-140Hz ($\gamma_{120-140}$) and 140-160Hz ($\gamma_{140-160}$). Average correlation values over subjects are displayed on a template brain where significant, as above.
